# Cave adaptation drives coordinated transcriptional remodeling across diverse cell types in the brain of a teleost fish

**DOI:** 10.64898/2026.06.19.733352

**Authors:** Edward S. Ricemeyer, Kathryn Gallman, X Maggs, Yulia Nussbaum, Rachel A. Carroll, Robert Peuß, Nicolas Rohner, Alex C. Keene, Wesley C. Warren

## Abstract

Adaptation of organisms to extreme environments requires dramatic phenotypic changes. Studying these changes can elucidate mechanisms underlying phenotypic differences in the context of both evolution and human disease. The Mexican tetra, Astyanax mexicanus, is a powerful model of extreme adaptation over a short evolutionary time scale. This fish species includes surface- and cave-dwelling ecotypes, with cavefish displaying many adaptations to subterranean life, including behavioral changes such as sleep loss, increased appetite, and reduced aggression. Unraveling the mechanisms underlying these changes has been challenging, presumably because they are complex traits that required coordinated changes across multiple cell types to evolve. Here, we present a spatially integrated comparative cell atlas of whole adult brains of surface and cavefish. After establishing the molecular signatures of 35 cell types, we show that cave colonization drove canalized regulatory changes to gene expression across diverse cell types. Cavefish brains show shifts in cell-type composition compared to their surface counterparts, as well as complex regulatory changes to pathways governing hypoxia response and circadian rhythm. Microglia in the cavefish brain underwent extensive transcriptional remodelling, including changes in senescence and AMPK pathways. Further, cell-cell communication analysis identified a cave-enriched ligand-receptor communication pattern centered on signals sent from glial cells to diverse populations of neurons. This atlas identifies genetic changes associated with neural and behavioral evolution and provides a resource for mechanistic studies examining brain evolution.

## Introduction

Adaptation to new environments is a major driver of biodiversity (Schluter 2000), but the genomic underpinnings of these changes remain poorly resolved. Thus, one of the central questions in evolution is how genomes adapt when faced with conditions different from those in which they evolved. Advances in technologies such as long-read sequencing (Rhoads and Au 2015) and genome assembly algorithms (Cheng et al. 2021) have made it straightforward and affordable to assemble and compare the genomes of closely related organisms adapted to extremely different environments (Guiglielmoni et al. 2024). However, identification of underlying mechanisms has remained elusive, in part because much of the change driving adaptation over short evolutionary time is based on regulatory changes rather than modifications to coding sequence (Wray et al. 2003). The ability to measure gene expression genome-wide using RNA-seq has been useful for finding genes differentially regulated between genomes adapted to different environments (Pavey et al. 2010). However, interpretation of these data can be difficult when analyzing a large number of complex traits that evolved via coordinated changes to the regulatory programs of multiple cell types (Zeisel et al. 2015).

Many examples of organisms displaying remarkable adaptations to extreme environments can be found in caves (Soares and Niemiller 2020). In the absence of light, brain structure and sensory processing display widespread, multi-level changes, acting through sensory, circadian, neuroendocrine, metabolic, and immune pathways (Soares and Niemiller 2020). Aside from perpetual darkness, these environments are also characterized by low oxygen (hypoxia), as well as changes to food availability and parasite load (Niemiller and Soares 2015). Many clades of animals contain species that have colonized subterranean environments and display common themes of sensory and behavioral adaptation compared to their aboveground relatives (Soares and Niemiller 2020). For example, the naked mole-rat *Heterocephalus glaber* has degraded eyes (Nikitina et al. 2004) and reduced neural visual processing pathways (Crish et al. 2006) compared to other rodents, but compensatory increases in tactile sensory sensitivity and pain tolerance (Vice et al. 2021). Similarly, the blind cave salamander *Proteus anguinus* has reduced eyes and visual nerves (Hawes 1945), but can navigate using magnetic fields (Roth and Schlegel 1988). Thus, across subterranean lineages, a reallocation of energetic and developmental resources moved investment away from the hardware and neural circuitry of vision and towards enhancements of non-visual sensory systems that help detect movement, vibration, smell, or electric fields. These perturbed processes often mimic human disease phenotypes, but their molecular signatures remain ill-defined.

A striking model of broader trait adaptation, including brain morphology, to extreme conditions can be found in the Mexican tetra, *Astyanax mexicanus* (Keene et al. 2016). This single species of fish comprises populations in the rivers of the Sierra del Abra region of northeastern Mexico in addition to at least 35 populations independently adapted to life in the surrounding caves (Mitchell et al. 1977). *A. mexicanus* cave evolution has occurred multiple times over the last approximately 20,000 years and is ongoing (Moran et al. 2023), with continuous hybridization between surface and cave populations. This provides a natural experiment for studying how rapid adaptation to a new environment drives evolution.

The *A. mexicanus* cavefish displays extreme phenotypic adaptations to the subterranean environment, including eye loss (Protas et al. 2007; Krishnan and Rohner 2017), starvation resistance (Aspiras et al. 2015), and enhanced olfaction (Bibliowicz et al. 2013; Espinasa et al. 2014). Some of the most intriguing, but least understood, differences between surface and cavefish lie in their behaviors (Kowalko 2020). Cave populations have lost their aggression (Burchards et al. 1985), schooling behavior (Kowalko et al. 2013), circadian rhythm (Duboué et al. 2011; Beale et al. 2013; North et al. 2025), and prey response (Yoshizawa et al. 2010). Previous studies have revealed the mechanisms underlying some of these behavioral changes, such as increased numbers of superficial neuromasts responsible for vibration attraction (Yoshizawa et al. 2010) and a mutation in *mao* resulting in reduced aggression (Elipot et al. 2013; Elipot et al. 2014). However, brains are highly heterogeneous, and most behavioral changes are likely polygenic, so identifying the genetic changes causing these phenotypes has been largely intractable. Single-cell and spatial RNA-sequencing present opportunities to break these changes down into components by cell type and brain region.

The brain contains many different cell types. These cell types have diverse functions ranging from the direct modulation of behavior performed by neurons to the immune surveillance provided by microglia (Prinz et al. 2019). Dysfunction of any of these cell types can lead to a wide-ranging set of diseases; for example, microglial dysfunction plays a role in human neurodegenerative diseases such as Alzheimer’s and Parkinson’s (Gao et al. 2023). Previous work has already shown major differences between the immune systems of surface and cavefish outside of the brain (Peuß et al. 2020), but immunity within the brain has not yet been studied. Thus, a central question is whether cell types in the brain evolved similarly during the process of adaptation to caves, or whether this change of environment put distinct evolutionary pressures on different cell types.

To gain insight into the complexities of cellular and molecular changes occurring in the cave-adapted brain, we present here a comparative cell-resolution atlas of the brains of surface- and cave-adapted Mexican tetras, integrating single-nucleus and spatial gene expression datasets. We use this atlas to explore differences in cellular composition, gene expression, signaling pathways, and cell-to-cell communication between surface and cave brains. We further investigate how these cellular differences relate to the adaptive changes in behavior and immunity present in cavefish. We expect this atlas, like previous cell atlases of whole fish brains (Hegarty et al. 2024) or brain regions (Shafer et al. 2022), to be highly valuable for future mechanistic studies of evolution of brain function in this and other species. Moreover, our results offer, for the first time, cell-level insight into how the brain adapts when faced with extreme environmental change.

## Results

### A spatially integrated comparative brain atlas

We present here a spatially integrated comparative atlas of the brains of adult surface and cave ecotypes of *A. mexicanus* (**Figure 1**). This atlas is composed of nuclei from the brains of seven Rio Choy surface fish (three males, four females) and nine Pachón cavefish (four males, five females), and spatial data from one female Rio Choy surface fish. After quality control, we used an unsupervised, network-based clustering process to partition 117k nuclei into 35 clusters (**Figure 1a**, **Supplementary Figure 1**). Integrating the spatial and nuclei data allowed us to resolve the location of each cluster in the brain (**Figure 1b-d**, **Supplementary Table 1, Supplementary Figure 2**). We assigned cell type labels to each cluster based on the canonical marker genes expressed in that cluster (**Supplementary Figure 3**, **Supplementary Tables 1-2**) and the spatial distribution in the brain of expression patterns associated with that cluster (**Supplementary Figure 2**, **Supplementary Table 1**).

**Figure 1.**
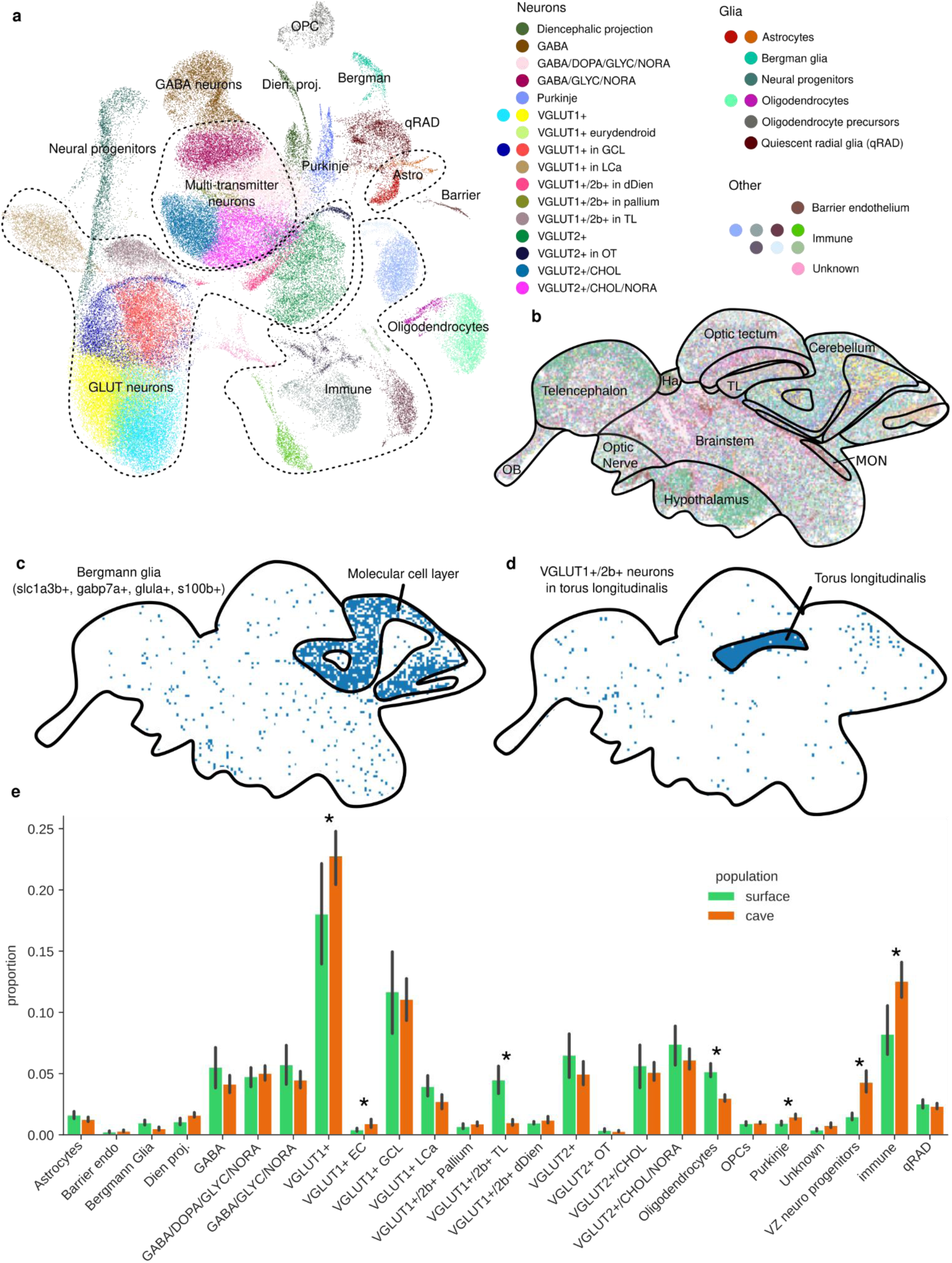
A spatially integrated comparative brain atlas. a, A UMAP representation of 117k nuclei from 7 Rio Choy surface and 9 Pachón cavefish, with cells colored by cell type and major groups outlined. **b,** A spatial map of a section of a surface fish brain, with spatial barcodes colored by their most similar cluster in the nuclei dataset. **c,** The same spatial map as **b**, but with only barcodes assigned to the Bergmann glia cluster, showing the concentration of this cell type expression pattern in the molecular cell layer of the cerebellum. **d,** The same spatial map as **b-c**, but with only barcodes assigned to the smallest of three clusters composed of glutamatergic neurons expressing both VGLUT1 and VGLUT2b. The spatial barcodes assigned to this cluster are concentrated in the torus longitudinalis, differentiating them from the other two VGLUT1+/2b+ clusters. **e**, Comparisons of proportions of each cell type in cave vs. surface brains. 95% confidence intervals shown. Stars indicate inferred differences at FDR = 0.05.

We then used the cell-type annotations to further refine the data. We merged clusters that could not be easily differentiated from each other by differentially expressed genes or spatial expression patterns, treating them as a single cluster in downstream analyses. For example, we merged clusters 0 and 1 into a single cell type, as both expressed genes indicative of glutamatergic transmission via vesicular glutamate transporter 1 (VGLUT1) and non-specific brain-wide spatial expression patterns. We also separated the immune cell clusters from the larger dataset. This allowed us to subset just the immune cells by cell type and delve deeper into comparative analysis of immune cell type diversity between surface and cave *A. mexicanus*.

Following refinement, we identified five top-level groups of neurons (**Figure 1a**, **Supplementary Figure 3a**): 1) nine distinct groups of glutamatergic neurons, broken down by spatial location and expression of vesicular glutamate transporters (e.g., **Figure 1d**); 2) four clusters of neurons each expressing transporters for multiple neurotransmitters; 3) a single cluster of purely GABAergic neurons; 4) a cluster of diencephalic projection neurons; and 5) a cluster of Purkinje neurons. We also found three top-level groups of glia (**Figure 1a**, **Supplementary Figure 3b**): 1) oligodendrocytes and their precursor cells; 2) quiescent radial glia, astrocytes, and Bergmann glia; and 3) a cluster of cerebellar ventricular zone neurogenic progenitors. Finally, we identified a cluster of barrier endothelium cells and assigned all immune cell types to a single category for subsequent higher-resolution reclustering.

### Cave adaptation accompanied by changes to neurons and glia

Comparative anatomical analysis has revealed differences in the size of brain regions between surface and cave populations of *A. mexicanus.* This includes an enlarged hypothalamus (Elipot et al. 2013; Elipot et al. 2014; Loomis et al. 2019) and habenula (Loomis et al. 2019), as well as a reduced optic tectum in cavefish (Sligar and Voneida 1976; Voneida and Sligar 1976; Soares et al. 2004; Loomis et al. 2019). Differences in the relative sizes of defined brain regions can indicate an increase in a particular population of neurons, such as the increase in serotonergic neurons in the hypothalamus of the cavefish (Elipot et al. 2013; Elipot et al. 2014), or a loss of sensory signal innervation, such as the reduction in the size of the optic tectum associated with the loss of visual system innervation in the blind cavefish (Loomis et al. 2019). In this way, comparative differences in brain region size can be indicative of locations where the nervous system of *A. mexicanus* evolved to facilitate cave life.

To investigate possible differences in cellular composition between surface and cavefish brains, we applied a Bayesian model for differential composition, with population and sex as covariates (**Figure 1e**, **Supplementary Figure 4**, **Supplementary Table 3**). We found that surface brains contain proportionally more oligodendrocytes than cave brains, as well as more VGLUT1+/VGLUT2b+ neurons located in the torus longitudinalis (TL). Cavefish brains contain proportionally more cerebellar ventricular-zone neurogenic progenitor cells, Purkinje cells, immune cells, VGLUT1+ eurydendroid neurons, and unlocalized VGLUT1+ neurons (FDR = 0.05).

### Cave adaptation is a major driver of differences in gene expression

In addition to differences in cellular composition between surface and cavefish brains, the cell atlas we present here allows for the examination of differences in gene expression at the cell-type level. To understand the role that ecotype plays in overall expression patterns in the brain, we divided cells into groups of cells from the same surface or cavefish brain sample and from the same cell type. We used these groups to perform hierarchical clustering with the UPGMA algorithm based on the first 20 principal components of expression across cell types. In addition to cell type and sample ecotype, we also considered the sex of each sample for comparison, because sex is known to be a major determinant of gene expression in the brains of other organisms (Parsch and Ellegren 2013). Qualitatively, cell type is the highest-level factor determining overall expression patterns, whereas ecotype and sex are both lower-level determining factors (**Figure 2a**).

**Figure 2.**
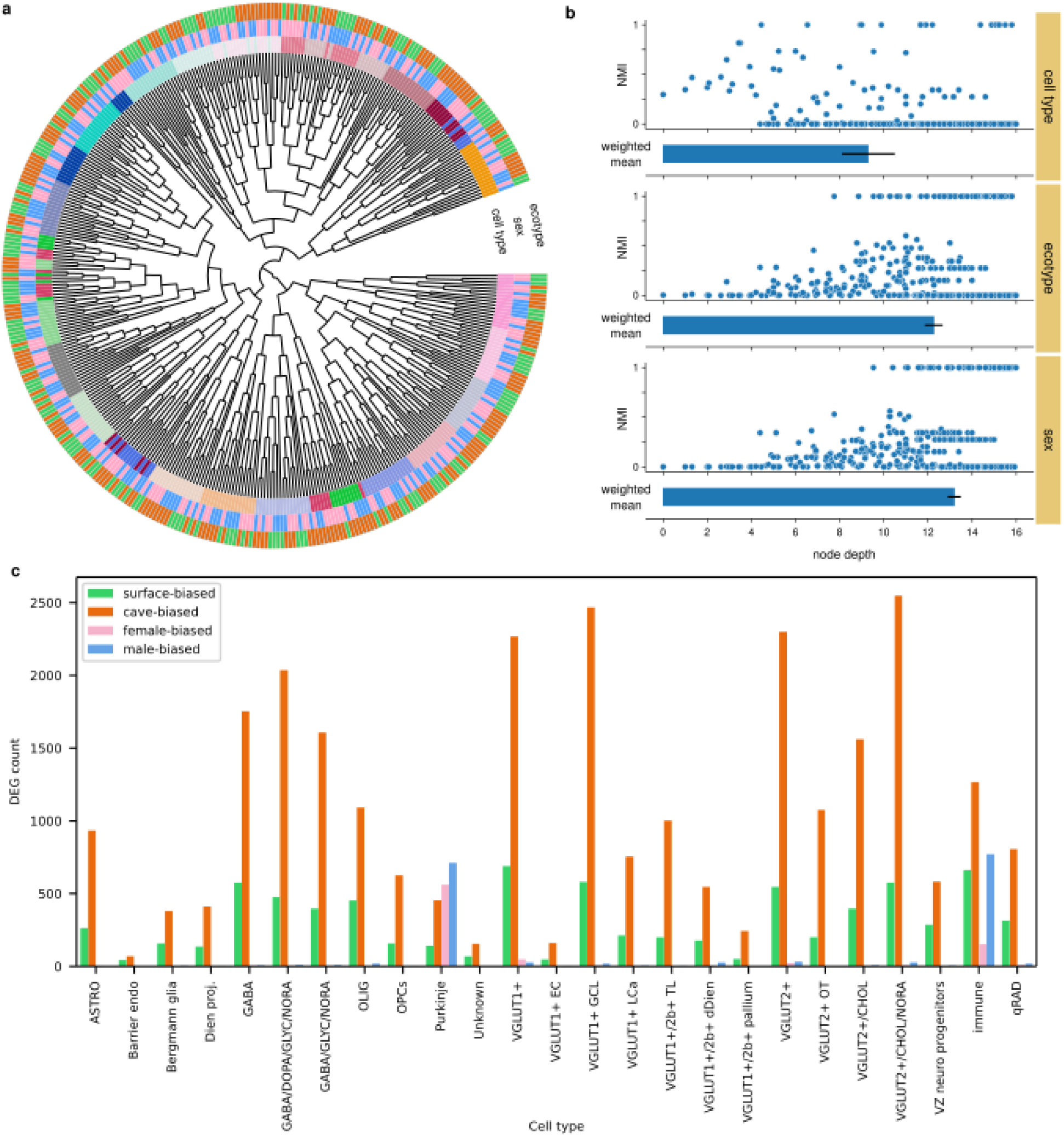
Cave vs. surface ecotype is the major driver of within-cell-type gene expression. **a**, Hierarchical clustering (UPGMA) based on gene expression of cells grouped by sample and cell type shows that cell type is the highest-level determinant of gene expression, with ecotype and sex as lower-level determinants. **b**, Normalized mutual information (NMI) versus node depth confirms that cell type is the highest-level determinant of gene expression, but also shows that ecotype is a higher-level determinant than sex. NMI is calculated for each inner node between descendant leaf character (i.e., cell type, ecotype, or sex) and leaf position (i.e., left or right descendant of inner node). Mean node depth weighted by NMI shown below each plot with bootstrapped 95% confidence intervals. **c**, Counts of DEGs per cell type for both cave vs. surface and male vs. female. With the exception of Purkinje cells, all cell types have more cave-biased genes than surface-biased or sex-biased genes.

To quantify this, we computed the normalized mutual information (NMI) between tree topology and character at each internal node of the tree for the three factors, where character is an ordinal encoding of cell type, ecotype, or sex. We then used this along with each node’s depth (that is, distance from the root of the tree) to calculate NMI-weighted mean depth for each factor, based on the insight that a higher NMI closer to the root for a given factor indicates that this factor is associated with higher-level organization of expression (**Figure 2b**). As expected, cell type is the highest-level factor, with NMI-weighted mean depth of 9.3 (bootstrapped 95% CI: 8.2-10.6). Ecotype is the second-highest-level determinant of expression tree topology, with NMI-weighted mean depth of 12.3 (95% CI: 11.9-12.7). Sex is the lowest-level factor, with NMI-weighted mean depth of 13.3 (13.0-13.6).

To gain a more concrete picture of how gene expression differs between surface and cavefish, we computed differentially expressed genes (DEGs) between surface and cave samples for each cell type in both the full dataset and immune subclustering datasets, using sample sex as a covariate (**Figure 2c**, **Supplementary Tables 4-5**). All cell types have at least some DEGs, with more DEGs overexpressed than underexpressed in cave compared to surface. VGLUT2+/CHOL/NORA, VGLUT1+ GCL, unlocalized VGLUT1+, and unlocalized VGLUT2+ neurons have the largest numbers of DEGs, although they are also the cell types with the largest numbers of cells, so this is likely to be, at least in part, an artefact of differing statistical power to detect DEGs depending on number of cells sequenced from each cell type.

Because of natural sample-to-sample variation and the large number of genes being tested, some differentially expressed genes may be found even between random groupings of samples. Therefore, to contextualize the amount of differential gene expression between surface and cave, we compared it to differential expression by sex, and between a random grouping of samples performed as a control. Accordingly, we found many genes with differential expression between males and females, although in agreement with the hierarchical clustering-based approach, there are fewer sex-biased genes than ecotype-biased genes (**Figure 2c**). Finally, to compare to the null hypothesis that ecotype drives differential gene expression no more than random chance, we randomly divided individuals into two equally sized groups regardless of sex or ecotype and ran DEG analysis again on each cell type with this arbitrary grouping as the only covariate. This resulted in a total of only three DEGs across all cell types.

Based on these analyses, we conclude that cave vs. surface is the major determinant of intra-cell-type variation in gene expression in the brain, more important even than sex.

### Cell-type specific differences in circadian-regulated genes

To better understand how these differences in gene expression relate to adaptations to subterranean life, we next performed enrichment analysis using Fisher’s exact test to find signaling pathways enriched for expression differences between surface and cavefish, focussing our interpretation on biological pathways linked to previously described cavefish traits, such as sleep loss and circadian rhythm dysregulation. Consistent with systemwide differences, a number of pathways were differentially regulated in many cell types, including circadian rhythm signaling (10 cell types), serotonin receptor signaling (9 cell types), and synaptic long term potentiation (9 cell types) (**Supplementary Table 6**), the latter suggesting cavefish neuronal remodeling has occurred. We also found differentially regulated pathways in specific cell types, such the hypoxia-inducible factor 1 (HIF1) pathway that is enriched in cavefish astrocytes compared to surface fish; we discuss this further in a subsequent section. These results indicate that both system- and cellular-level changes in expression of key molecular pathways were induced through adaptation to the cave environment in *A. mexicanus*.

Cave environments are largely arrhythmic, with perpetual darkness and limited fluctuations in temperature. Thus, the regulation of normally rhythmic behaviors and biological processes is a key hurdle for cave adaptation. Bulk RNA-seq has previously measured body-wide changes in the transcription of genes involved in regulating circadian rhythms between surface and cave ecotypes of *A. mexicanus*, with a loss of rhythmicity in many genes within the core circadian transcriptional translational feedback loop (TTFL) (Mack et al. 2021; Olsen et al. 2023). However, the cellular basis of these changes remains largely unresolved. We observed differential expression of subsets of *A. mexicanus* orthologs of mammalian core clock component proteins BMAL, CLOCK, CRY, and PER across all cell types. No single gene was differentially expressed in every cell type, highlighting the importance of measuring changes to gene expression at the cell-type level (**Figure 3a-b**). At the bulk RNA-seq level, cryptochrome genes *cry1a*, *cry1b*, *cry3a*, and *cry3b*, which all act as repressors in the core circadian TTFL (Liu et al. 2015), are rhythmic in surface fish while all except *cry1b* lose their rhythmicity in Pachón cavefish (Mack et al. 2021). We found that all of these genes are differentially expressed in varied but overlapping subsets of cell types in the brain; *cry1a* and *cry3a* had higher expression in cavefish, but *cry1b* and *cry3b* had higher expression in surface fish (**Figure 3b**). Conversely, the period circadian proteins *per1a*, *per1b*, and *per3*, which also serve as repressors to circadian gene transcription (Dunlap 1999), maintain their rhythmicity in Pachón (Mack et al. 2021), with increased expression in many cell types compared to surface fish brains (**Figure 3b**).

**Figure 3.**
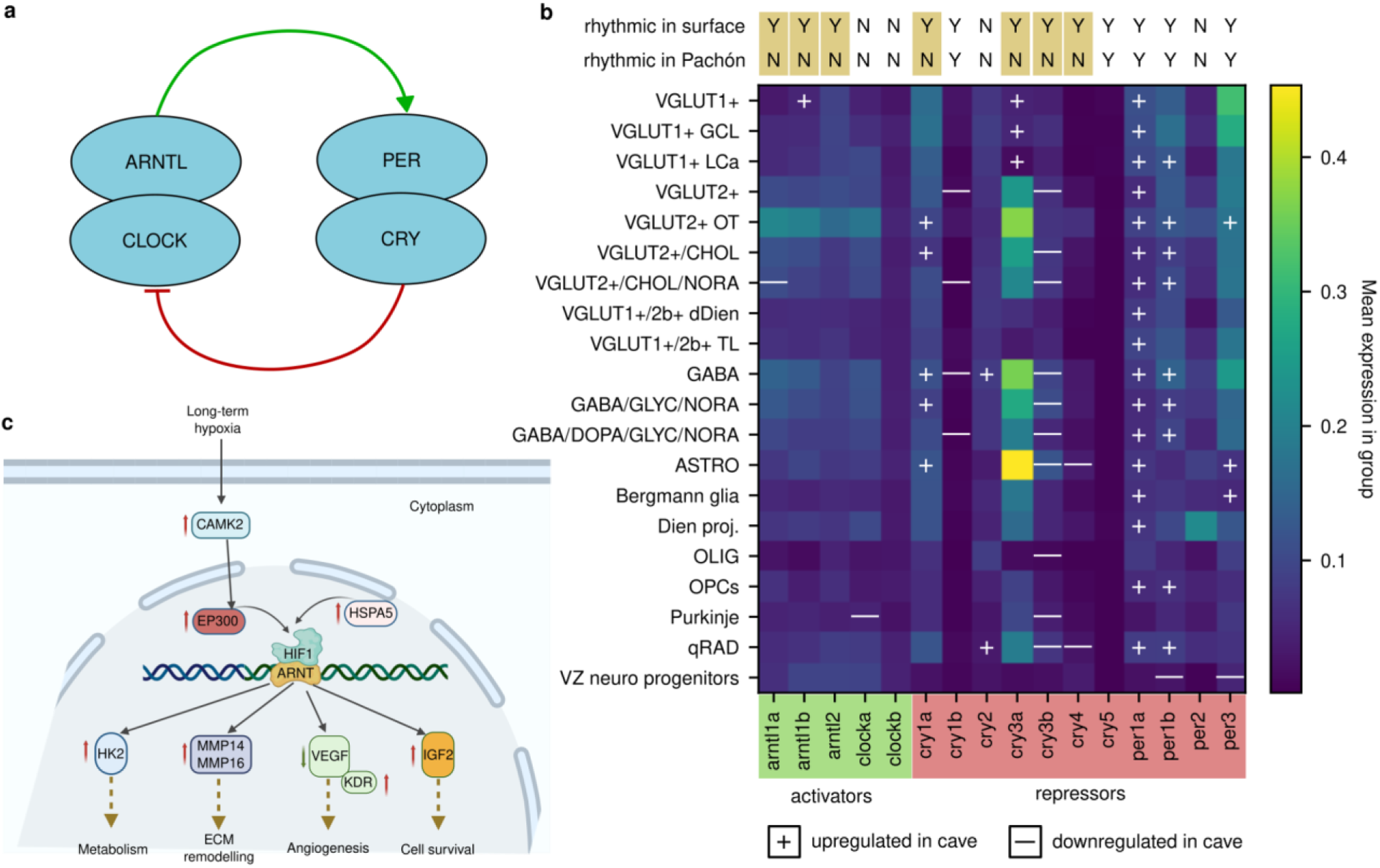
Cavefish have evolved cell-type specific changes to circadian rhythm and hypoxia response pathways. **a**, Circadian rhythm is controlled by two heterodimers in a negative feedback loop: ARNTL/CLOCK and PER/CRY. Previous work demonstrates that in surface fish, many of the genes coding for components of this feedback loop are expressed rhythmically over the course of a day, but in cavefish, most of these genes lose their rhythmicity. **b**, Mean and differential expression of core circadian rhythm complex genes by cell type. Many of these genes are expressed at different levels in cavefish compared to surface fish in different cell types, but there is no clear link between differential expression and loss of rhythmicity or the role of a gene in the feedback loop (i.e., activator vs. repressor). **c**, Changes to HIF1 signaling alter expression in astrocytes of genes involved in glucose metabolism, extracellular matrix (ECM) remodelling, angiogenesis, and cell survival.

Different cave lineages of *A. mexicanus* cavefish have evolutionarily converged towards similar phenotypes. In many cases the mechanistic changes that lead to these phenotypes occur through different points in the same physiological pathway. This is likely the case for the *per3* genes in *A. mexicanus* cave phenotypes (**Figure 3b**). Our data show an increase in transcription of *per3* in cavefish from the Pachón cave, but *per3* expression is reduced in the Molino cave ecotype of *A. mexicanus* through a high-frequency heterozygous deletion of its canonical start codon (Warren et al. 2021). This opposite direction of expression regulation in different cave morphs of the same gene, accomplished through coding sequence deletion in Molino but a regulatory change in Pachón, underscores the multiple routes through which different lineages of *A. mexicanus* have perturbed the core circadian TTFL to disrupt circadian rhythm. In other cases, the type of genetic modification for the evolution of a particular cave phenotype is more tightly conserved. We find an increased expression of *cry1a* in Pachón cavefish compared to surface fish (**Figure 3b**), a gene known to have lost rhythmic expression in this cave ecotype (Mack et al. 2021). The mutation reported in *cry1a* in the Pachón cave ecotype of *A. mexicanus* (Moran et al. 2022) has also been reported to occur as a substitution of the same amino acid in Somalian cavefish as well as in subterranean mammals, such as the naked mole rat (Moran et al. 2023; Swaminathan et al. 2025). This repeated evolution of circadian clock genes suggest an adaptive advantage to circadian dysregulation in the subterranean environment. *Cry1a* in particular is directly photoentrained by light in fishes and increasing expression consistently rather than rhythmically may allow for the continual regulation of downstream biological processes dependent on *cry1a* expression.

Activators of the core circadian TTFL are also differentially expressed across cell types. *Clocka* (Dunlap 1999) is arhythmic in both surface and cavefish (Mack et al. 2021), but only shows a change of expression in Purkinje cells, where it is reduced in cavefish (**Figure 3b**). *Arntl1a* and *arntl1b* both lose their rhythmicity in Pachón (Mack et al. 2021), but *arntl1a* is downregulated in cavefish VGLUT2+/CHOL/NORA neurons whereas *arntl1b* is upregulated in cavefish VGLUT1+ neurons (**Figure 3b**), suggesting that these two orthologs of mammalian ARNTL1 may have taken on different roles or at least were targeted in opposite direction by selection after cave colonization. Curiously, knockdown (KD) of ARNTL1 in the suprachiasmatic nucleus (SCN) of mice causes behavioral changes including anxiety-like behavior and increased stress sensitivity (Landgraf et al. 2016). When ARNTL1-KD mice are kept in total darkness, there is an extension of the free-running period of locomotor activity, typically entrained by light/dark cycles (Landgraf et al. 2016). Thus, the variable expression of *arntl1a* and *arntl1b* across different neuronal cell types in the brain may speak to cell-specific trade-offs regarding the impacts of ARNTL1 on stress and locomotor behavior, influencing the increased locomotor activity and decreased stress responses of cavefish. We find no clear pattern relating differential expression of these genes between surface and cavefish to each gene’s role as an activator or repressor within the core circadian TTFL, or to whether a gene lost its rhythmicity during cave adaptation. This, along with the different sets of cells displaying differential expression of each gene involved and the presence of cave-specific mutations in multiple genes in this feedback loop, demonstrates that adaptation of circadian rhythm to the arhythmic cave environment likely involves coordinated multigenic regulatory changes across different cell types. Time-course single-cell experiments with repeated sampling over the course of a day, as well as additional genomics data beyond RNA-seq, will be necessary to further disentangle the roles of different cell types in this adaptation. Nonetheless, this work is the first demonstration of the feasibility of breaking down expression of circadian rhythm genes by cell type in the cavefish brain.

### Canalized hypoxia signaling in cavefish astrocytes

The lack of sunlight in cave environments also prevents the growth of organisms that release oxygen through photosynthesis, making closed cave systems relatively hypoxic (Boggs and Gross 2021). In addition to differences in the expression of circadian rhythm pathway genes across cell types, we also discovered differences in the regulation of hypoxia-inducible factor 1 (HIF1) pathway in astrocytes (**Figure 3c**). HIF1A is a transcription factor that under normoxic (normal oxygen levels) conditions is hydroxylated and degraded, but under hypoxic (low oxygen) conditions stabilizes and promotes gene expression patterns that mitigate low oxygen availability (Majmundar et al. 2010). Cavefish adapted to life in hypoxic subterranean environments (Boggs and Gross 2021) and upregulate expression of HIF1A orthologs *hif1aa* and *hif1al2* and their targets during development, even under normoxic conditions (van der Weele and Jeffery 2022). Downstream effects of this overexpression, especially via the HIF1 pathway’s upregulation of sonic hedgehog signaling, are hypothesized to include both adaptations to the cave environment such as enhanced olfaction (Yamamoto et al. 2004) and taste bud amplification (Yamamoto et al. 2009) as well as side effects without an obvious adaptive advantage such as heart asymmetry reversal (Ng et al. 2026).

We found 16 genes in the HIF1 signaling pathway in astrocytes with differential expression between surface and cavefish (**Supplementary Table 6**). The gene in this pathway with the most extreme expression difference was *slc2a1*, with a much higher expression in cavefish (LFC = 3.23; multiple hypothesis corrected p = 4.27 × 10^-7^); this gene’s human ortholog is a direct target of HIF1A (Abu El Maaty et al. 2022). Other *A. mexicanus* orthologs of HIF1A signaling targets with upregulated expression in cavefish astrocytes include *hk2* (LFC = 1.22; p = 5.68 × 10^-3^), which codes for a glucose phosphorylation enzyme necessary for the first step of glucose metabolism and also is activated during starvation to maintain homeostasis (Tan and Miyamoto 2015); *mmp14b* (LFC = 1.44; p = 7.46 × 10^-3^) and *mmp16b* (LFC=2.46; p=8.43 × 10^-3^), which code for proteins involved in extracellular matrix remodelling (Sato et al. 1994); and *igf2b* (LFC = 1.22; p = 1.13 × 10^-3^), coding for an insulin-like growth factor. Conversely, *vegfc* (LFC = -1.44; p = 5.90 × 10^-5^), which promotes angiogenesis (Oh et al. 1997), has downregulated expression in cavefish. Given the roles of astrocytes in maintaining the blood-brain barrier, regulating nervous system homeostasis, protection against oxidative stress, and shaping connectivity between brain regions (Cooper et al. 2026), these hypoxia signalling-dependent changes in gene expression in astrocytes are likely to have far-reaching effects on brain function. Untangling the phenotypic effects of this large perturbation to the gene expression of astrocytes will require more specific experiments, but this is the first evidence implicating evolution of astrocytes in cave evolution, and we expect it to be a fruitful line of inquiry based on the many other previously established ways changes in HIF1 signalling are responsible for drastic cave-associated phenotypes (Yamamoto et al. 2004; Yamamoto et al. 2009; van der Weele and Jeffery 2022; Ng et al. 2026).

### Immune subclustering identifies diverse repertoire, with differences between surface and cavefish

The clustering resolution that best divided the full set of nuclei into biologically relevant and distinguishable groups of neurons and glia was too low to resolve immune cell subtypes, so we reclustered the subset of all nuclei assigned to immune clusters, starting with their original raw counts. As with the full set of nuclei, we assigned labels to each cluster based on expression of canonical marker genes (**Figure 4a**, **Supplementary Tables 7-8**, **Supplementary Figures 5-6**). Immune cell types did not show spatial patterns, so we were not able to add spatial annotations to these clusters (**Supplementary Figure 2**). We found a full immune repertoire comprising B cells, T cells (including gamma-delta and double-positive clusters), macrophages, microglia and their precursors, basophils and mast cells, and neutrophils.

**Figure 4.**
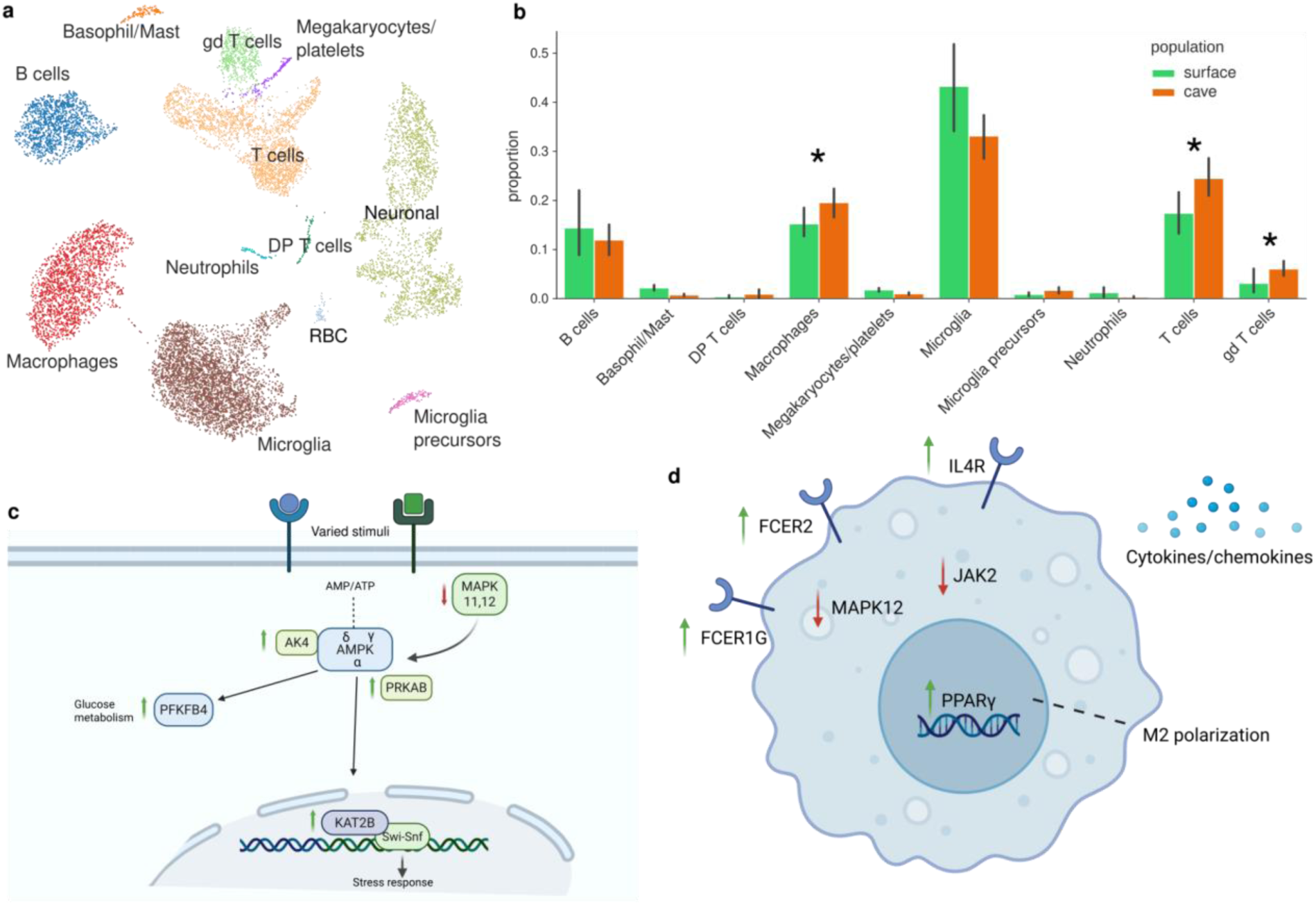
Immune cell types in the *A. mexicanus* brain. **a**, Cell types identified in the immune subclustering. DP T cells: double positive T cells. gd T cells: gamma delta T cells. RBC: red blood cells. **b**, Proportional changes in immune cell types when comparing cave to surface fish. 95% confidence intervals shown. Stars indicate inferred differences at FDR = 0.05. **c**, AMPK signaling pathway in brain microglia. AMPK (AMP-activated protein kinase) is a master metabolic sensor that maintains cellular energy homeostasis, activated by high AMP:ATP ratios caused by fasting (low energy sources in the cave environment). Our findings suggest upregulation of the AMPK pathway in cavefish brain microglia. **d**, Alternate macrophage activation pathway that indicates a M2 anti-inflammatory subtype in the brain.

Comparing compositions of immune cell types between surface and cavefish, we found that cavefish have proportionally more T cells, gamma delta T cells, and macrophages (**Figure 4b**, **Supplementary Table 9**; FDR = 0.05) than surface fish. This is consistent with the longer and more sensitive pro-inflammatory cytokine response and increased investment of the adaptive immune system previously described outside of the brain in cavefish (Peuß et al. 2020). We also found sex differences in composition: male brains contained proportionally more microglia and fewer B cells than female brains (**Supplementary Figure 7**). An increased number of microglia in male brains is consistent with evidence that microglia are critical contributors of brain sex differences in mammalian brains (Lenz and McCarthy 2015) where sex-specific differences in the number of microglia across males and female brains ontologically varies in a region-specific manner starting in early development (Han et al. 2021).

### Differential regulation of immune pathways

Cave adaptation in *A. mexicanus* involved broad changes to the immune system, with the lower parasite diversity in the subterranean environment leading to shifts from investment in innate immunity towards adaptive immunity (Peuß et al. 2020). Reductions in overall composition of immune cell types in the hypothalamus of cave versus surface fish were previously reported (Shafer et al. 2022), but little is known about the molecular phenotypes of immune cells in *A. mexicanus* whole brains, or how they differ between surface and cavefish. We looked for gene pathways with enrichment for differential expression between surface and cavefish in immune cell types by performing DEG analysis separately on each cluster (**Supplementary Tables 10-11**). We confined our analysis to microglia, macrophages, and T cells because these were the only groups with sufficient numbers of DEGs to find pathway enrichment, with DEG counts in the hundreds. As with neuronal and glial cell types, each immune cell type had more genes with higher expression in cavefish than in surface fish, demonstrating system-wide remodeling of regulatory networks in the brains of cavefish where increased expression supports canalization of predominately regressive brain-associated phenotypes.

Sleep deprivation is known to reduce microglial function and thus impair synaptic pruning in other organisms (Tuan and Lee 2019), which raises the question of whether cavefish, given their reduced sleep (North et al. 2025), have avoided this impairment through alterations to microglia. We found 521 genes with at least twofold differential expression (p < 0.05) between surface and cavefish microglia, the most of any immune cell type in the *A. mexicanus* brain. Although comparisons of the numbers of observed DEGs are susceptible to biases, this large number of DEGs nonetheless suggests the possibility that microglial function is altered in cavefish. Cell senescence is the top-scoring enriched pathway for these DEGs (-log BH p-value = 3.44), with several orthologs of transcription factors upregulated in cavefish microglia: *chp1*, *phf1*, and *smad1* (**Supplementary Table 12**). The mouse ortholog of cavefish *smad1* suppresses cell senescence by inhibiting expression of senescence-related genes (Xu et al. 2022), and the human ortholog of *phf1* is a positive regulator of the p53 pathway (Yang et al. 2013). Thus, the transcription factor expression profiles reflect an evolutionary adaptation toward non-inflammatory tissue maintenance, with evidence that microglia have adapted to be more resilient in the face of sleep deprivation.

Another notable enriched pathway in cavefish brain resident microglia is the AMPK pathway, known to be activated by low glucose (Mihaylova and Shaw 2011) (**Figure 4c**). Because we sequenced fish that were reared in a laboratory environment, the activation of this pathway in cavefish, despite good food availability, may be a demonstration of canalization of a response to low glucose that now persists regardless of actual glucose levels, which are generally higher in cavefish due to their insulin resistance (Riddle et al. 2018). A -2.27 z-score is the result of 12 of 110 genes that have measurement direction consistent with a decrease in movement disorders.

Macrophages are normally excluded from the brain, but are sometimes able to cross the blood-brain barrier as a result of injury, inflammation, infection, or developmental remodeling (Ransohoff and Engelhardt 2012). We found macrophages in both surface and cavefish brains, which we also evaluated for pathway enrichment based on the 324 DEGs between the two ecotypes. Of the 17 enriched pathways, several suggest altered immune surveillance and inflammatory signaling (**Supplementary Table 12**). Another enriched pathway, macrophage alternative activation pathway (**Figure 4d**), shows upregulation of FcERI resulting in SYK activation, a kinase strongly implicated in microglia-mediated synapse removal (Li and Barres 2018).

For T cells, we observe 34 enriched pathways, with a majority categorized as related to cytokine signaling (**Supplementary Table 12**). A top-scoring pathway is FLT3 signaling (-log10 corrected p-value = 3.55). FLT3 is a regulator of T-cell progenitors and dendritic cells, and its expression in T cells usually denotes early progenitor-like states. Upstream regulators include a network associated with TCR activation, a process leading to increased expression of transcription factors such as JUNB and MYC, as well as their dimerization partner MAX. These factors play crucial roles in regulating gene expression programs necessary for T cell activation, proliferation, and differentiation, reflecting a direct regulatory cascade triggered by TCR engagement. Furthermore, increased expression of BTLA, SOCS2, and CFLAR following an assumed TCR stimulation suggests that their induction in the cavefish brain may be one mechanism of the previously observed shift of investment from the innate immune system towards homeostasis-promoting T cell populations (Peuß et al. 2020). Specifically, BTLA negatively regulates T cell activation, SOCS2 inhibits cytokine signaling, and CFLAR blocks apoptotic signaling (Sedy et al. 2005; Linossi et al. 2013). Further data will be needed to validate their state in surface and cavefish brains.

These results demonstrate the importance of including immune cells when characterizing differences between surface and cavefish brains. The extensive transcriptional remodeling we observe in microglia, macrophages, and T cells indicates that the immune system in the brain is highly divergent between the two ecotypes. The functional implications of these immune system adaptations extend beyond its traditional role of fighting infection, with microglia possibly playing a role in the synaptic pruning that reduces optic tectum size and overall alters brain neuronal connectivity in the cavefish (Tuan and Lee 2019; Mendez Scolari et al. 2026).

### Reconfigured astrocyte to neuron communication

Differences between surface and cavefish at the level of individual cell types yield many observations about differences in gene expression between surface and cavefish, and more broadly, how these differences affect various molecular interaction pathways within specific cell types. One recurring theme of these observations is that adaptation required coordinated changes to expression of different genes and pathways in different cell types. To better understand how such adaptations within each cell type interact to achieve phenotypic changes, we next looked for changes in communication between cell types using two complementary approaches for understanding differences in ligand-receptor (LR) interactions between surface fish and cavefish: tensor decomposition and DEG-based analysis.

We used tensor decomposition to deconvolute the full set of intercellular interactions across the brain into modules representing smaller communication patterns. This analysis found five factors, each representing a set of related LR interactions broken down by sample, sender cell type, receiver cell type, and specific pair of interacting genes (**Figure 5a**). Of these five patterns, one has a difference in activity between surface and cavefish: Factor 4 is enriched in cavefish compared to surface fish (**Figure 5b**). Interactions in factor 4 are characterized by astrocytes, Bergmann glia, and quiescent radial glia as the sender cells (that is, the cells expressing genes coding for ligands) and neurons as the receivers (that is, the cells expressing genes coding for receptors), hinting at a possible rebalancing of the brain’s intercellular signaling architecture that warrants further investigation. This could present as increases in metabolic support for neurons, homeostatic regulation, synaptic plasticity, and neuromodulation (De Pittà and Brunel 2016; Zabegalov et al. 2021; Pathak and Sriram 2023; Verkhratsky et al. 2026).

**Figure 5.**
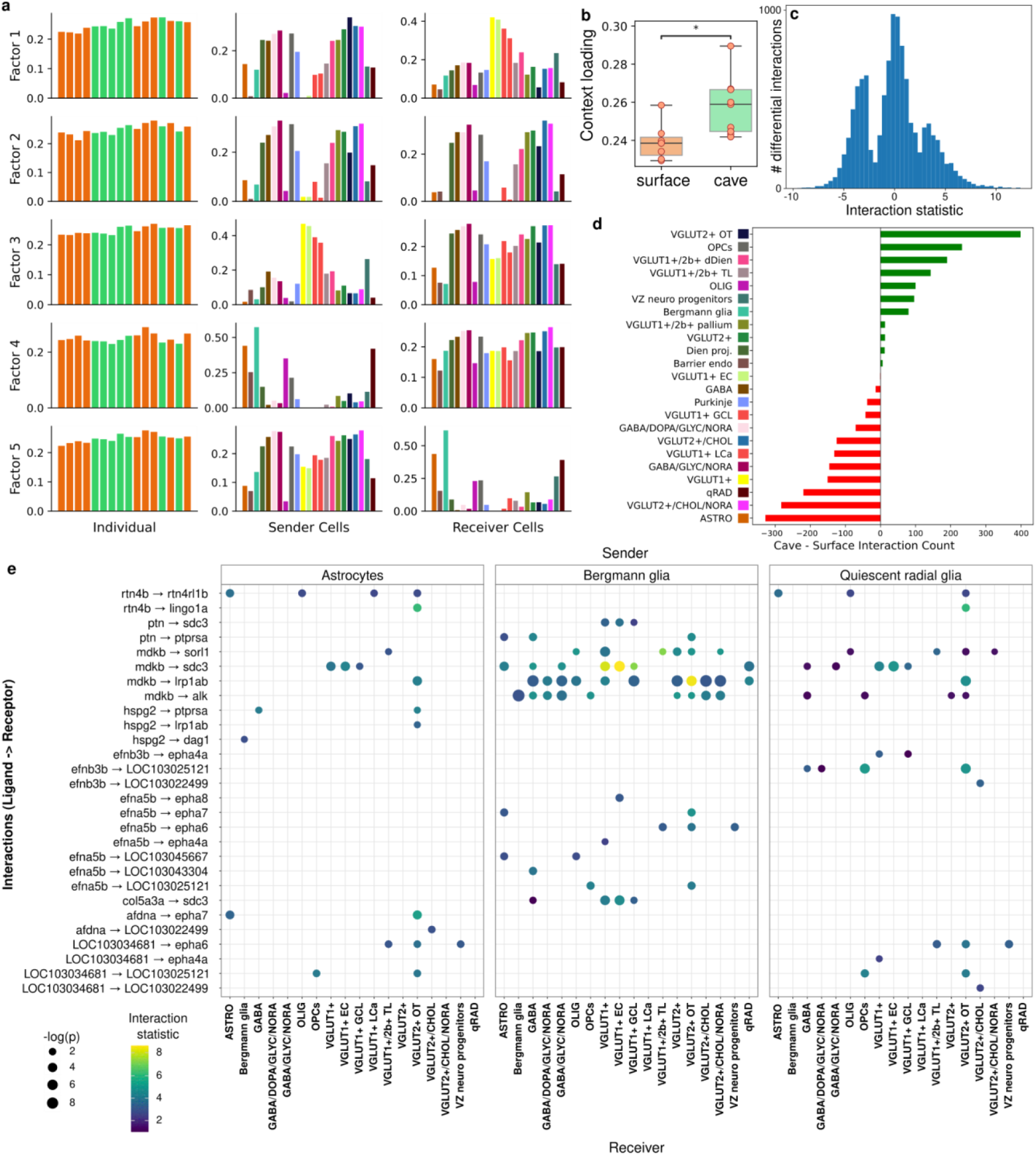
Cell–cell communication analysis between surface and cavefish. **a**, Normalized loadings (unit Euclidean length) for samples, sender cell types, and receiver cell types, showing relative importance of different variables in five cell-cell communication programs (factors). Bars are colored according to population or cell type. **b,** Sample loadings for surface and cavefish populations within factor 4. Boxes represent interquartile ranges, with whiskers indicating the full distribution. Groups were compared using a two-sided independent *t*-test followed by Bonferroni correction. *, *p* < 0.05. **c**, Interaction statistic distribution for all differentially expressed ligand-receptor interactions (*p* < 0.05) between surface and cavefish inferred from DEGs. Positive interaction statistics indicate enrichment in Pachón samples, whereas negative values indicate enrichment in surface fish. **d**, Relative involvement of cell types in surface- or cave-enriched ligand-receptor interactions. **e**, Ligand-receptor interactions identified by both Tensor-cell2cell (Factor 4) and DEG-based inference, restricted to interactions with astrocytes, Bergmann glia, and quiescent radial glia as sender cell types and all other cell types as receivers.

In parallel, we mapped known ligand-receptor interactions onto differentially expressed genes between the two populations, revealing 11,294 interactions that are differentially regulated between surface and cavefish (**Figure 5c**; BH-corrected p < 0.05). We also examined the contribution of individual cell types to ligand-receptor interactions in surface and cavefish. This analysis revealed that VGLUT2+ OT, OPCs, VGLUT1+/2b+ dDien, VGLUT1+/2b+ TL, oligodendrocytes, ventricular zone (VZ) neural progenitors, and Bergmann glia participated more frequently in ligand-receptor interactions in cavefish than in surface fish. In contrast, astrocytes, VGLUT2+/CHOL/NORA, quiescent radial glia, VGLUT1+, GABA/GLYC/NORA, VGLUT1+ LCa, and VGLUT2+/CHOL cell types showed greater involvement in surface fish interactions (**Figure 5d**).

To find overlap between the two independent analyses, we selected the top ligand-receptor interactions contributing to Factor 4 (values > mean + 2 SD; n = 81) and intersected them with differentially expressed interactions, resulting in 61 shared interactions. These interactions were visualized using astrocytes, Bergmann glia, and quiescent radial glia as sender cell types and all other cell types as receivers (**Figure 5e**). Several ligands recur repeatedly in these results, with enriched interactions with multiple receptors in many sender/receiver cell type pairs. For example, *mdkb*, an ortholog of mammalian MDK, which promotes neurite outgrowth and mediates signaling between the nervous system and the immune system (Neumaier et al. 2023), shows enriched interactions with four different receptors. Each of these four interactions is enriched across more than 10 glial sender to neuronal receiver cell-type pairs. This, along with the similar roles of astrocytes, Bergmann glia, and quiescent radial glia in teleost brains, suggests an astrocyte-led reconfiguration of neural connectivity and modulation of nervous system inflammation.

Together, these results show that substantial reorganization of intercellular signaling in the brain accompanied the transition to subterranean life. Relating these complex multifactorial modifications in communication patterns that accompanied adaptation to caves to specific phenotypic changes will require further experimentation, but our study provides a promising set of targets for this future work.

## Discussion

We present here the first single-cell-resolution, spatially integrated, comparative brain atlas of a vertebrate adapted to an extreme environment, providing a resource that substantially advances understanding of how neural circuits, glial physiology, and brain immunity evolve at the molecular level. By integrating single-cell and spatial data across the whole brain, we are able to move beyond the anatomical and bulk transcriptomic comparisons that have characterized most prior work on *A. mexicanus* brain evolution (Loomis et al. 2019; Jaggard et al. 2020; Kozol et al. 2023) and resolve the molecular basis of differences that were previously invisible at lower resolutions. Several themes emerge from this cell-level view. First, the magnitude and ubiquity of differences in expression between surface and cave ecotypes within a single species, even when raised in an identical laboratory environment, suggests that adaptation to the subterranean environment has led to modifications in transcriptional regulation throughout the brain, rather than merely in regions like the optic tectum, which is directly linked to the loss of visual cues. Second, the changes we observe are consistently distributed across multiple cell types rather than concentrated in any single population, reinforcing the view that cave-associated traits such as circadian rhythm disruption and altered immune function are the products of coordinated multigenic regulatory evolution (Wray et al. 2003; Zeisel et al. 2015). Third, and perhaps most strikingly, the immune system of the cavefish brain has undergone extensive remodeling that parallels, and may help explain, previously described immune differences between ecotypes (Peuß et al. 2020). Together, these findings highlight the power of single-cell approaches for dissecting the genomic basis of complex adaptive phenotypes and establish a foundation for future mechanistic discovery in this system.

In addition to enabling the biological insights we present here, this cell atlas will be a valuable resource for future studies of brain adaptation in *A. mexicanus*, an emerging model for both evolution and human diseases such as metabolic and neurodegenerative disorders. For example, bulk RNA-seq data can be deconvoluted using this atlas, providing cell-type labels without the need for single-cell experiments. Moreover, future single-cell datasets can be projected onto our atlas, avoiding the need for manual annotation. Finally, the atlas serves as a reference of gene expression levels between cell types and ecotypes that can be easily accessed through the UCSC cell browser (see data availability).

In this study, we predicted differences in gene interactions and signalling pathways and related these differences to known phenotypic changes associated with cave adaptation. Few pathway resources exist for *A. mexicanus*, or even teleosts in general, so our approach, commonly employed in studies of non-model organisms, required projecting expression data from *A. mexicanus* genes onto their mammalian orthologs to use pathway and interaction databases based on the primary mammalian model organisms, mice and humans. Many of the differentially regulated pathways we inferred recapitulate previously known differences between surface and cave brains, such as changes in sleep and hypoxia signaling, confirming the value of this orthology-based approach. However, there are limitations to this strategy of projecting gene expression onto pathways in a distantly related species based on protein sequence orthology. Most importantly, the common ancestor of teleosts underwent a whole-genome duplication event, resulting in *a* and *b* orthologs of many mammalian genes that often underwent subsequent neo- or subfunctionalization (Glasauer and Neuhauss 2014). Moreover, even in more closely related pairs of species not separated by a drastic event like whole genome duplication, such as humans and mice (Stergachis et al. 2014; Bravo González-Blas et al. 2023) or zebrafish and *A. mexicanus* (Shafer et al. 2022), primary regulators of cell fate are conserved but the genes and *cis*-regulatory elements they target have high turnover. Therefore, establishing lineage-specific cell-type-resolved gene regulatory networks in *A. mexicanus* will be necessary for future determination of the precise mechanisms through which evolved genetic changes between surface and cavefish lead to gene expression changes that in turn cause cave-specific behavioral phenotypes. This work will require collecting single-cell multi-omic data to find transcription factor binding sites and map chromatin structure and accessibility throughout the genome.

One striking example of this is our finding of differential HIF1A signaling in astrocytes between surface and cave ecotypes. Much previous work has established the outsized pleiotropic role of changes in HIF1A signaling, specifically the canalization of the hypoxia response, in producing many cave-associated phenotypes during development (Yamamoto et al. 2004; Yamamoto et al. 2009; van der Weele and Jeffery 2022; Ng et al. 2026), so changes to gene expression downstream of HIF1A in astrocytes may be at least partially causative of adaptations in the cavefish brain. Given the role of astrocytes in both circadian rhythm (Brancaccio et al. 2019) and nervous system repair (Anderson et al. 2016), the relationship between HIF1A signaling in astrocytes and adaptations such as sleep loss is worth investigating with subsequent experiments. A first step in this direction will be to establish the locations of HIF1A-targeted response elements in astrocytes, along with differences in sequence and function of these elements between ecotypes; this will require collecting single-cell ATAC-seq data. Then, perturbation of these regulatory sequences using CRISPR can be used to determine which changes are responsible for phenotypic differences between surface and cavefish, including the molecular phenotypic differences we describe in this study.

Ultimately, the data presented here demonstrate that adaptation to an extreme environment requires pervasive, cell-type-distributed remodeling of transcriptional programs across neurons, glia, and immune cells simultaneously. Without single-cell transcriptomics, reaching this conclusion would have required multiple large-scale studies of individual tissues and cell types. The *A. mexicanus* system is particularly well-suited to building on these findings, given the availability of multiple independently evolved cave populations, the possibility of generating surface-cave hybrids, and an increasingly complete toolkit of genomic and genetic resources, including a platform for large-scale CRISPR screening (Shennard et al. 2025), a catalog of structural variation (Roback et al. 2025), and recent cell atlases of the olfactory epithelium (Choi et al. 2026) and embryonic brain (Gallman et al. 2026). The differentially regulated pathways and cell-cell communication programs identified here thus provide prime targets for future studies of mechanisms of adaptation. Such studies could focus on determining which of the hypoxia signalling pathway changes may be causally linked to sleep loss, establishing whether the pro-senescence microglial state confers neuroprotection under chronic metabolic stress, and resolving the functional consequences of the reorganized glial-neural communication network. More broadly, several of the cave-associated molecular signatures identified here, such as microglial AMPK activation and hypoxia-dependent astrocyte reprogramming, mirror mammalian models of neurodegenerative disease (Zhou et al. 2022; Gao et al. 2023). The cavefish brain therefore represents a window into the evolutionary biology of extreme adaptation as well as a naturally occurring model in which disease-relevant transcriptional states have been canalized by selection, offering new opportunities to study their causes and consequences outside of a pathological context.

## Methods

### Sample collection

All *A. mexicanus* tissue samples were processed from fish reared at the Stowers Institute for Medical Research aquatic facility under IACUC approved protocol (Protocol ID: 2021-126). For single nuclei isolation, seven adult Rio Choy surface fish (three males, four females) and nine adult Pachón cavefish (four males, five females) were euthanised with hypothermal shock in an ice bath. All individuals were between 1-2 years old. Whole brains were harvested and immediately flash frozen in liquid nitrogen. On the day of dissection, we systematically rotated between dissecting cave and surface individuals to avoid bias in regulation of the circadian rhythm across ecotypes. Dissections took approximately 8 hours, over the course of two days, dissecting females in March 2023 and males in January 2024.

Nuclei were isolated from cryopreserved adult brain samples using the Singulator2 instrument per manufacturer’s protocols (S2 Genomics; Livermore, CA). After mechanical disruption, cell filter straining steps, and two washes at 250ul each, the cell nuclei were suspended in a nuclei isolation buffer (S2 Genomics; Livermore, CA), with all buffers containing 0.4U/ul Millipore RNAse inhibitor (SigmaAldrich). Nuclei were counted and evaluated for the amount of remaining cellular debris using the Cellometer instrument (Revvity; Waltham, MA).

For the spatial experiment, one adult female was euthanised with hypothermal shock. Whole brain tissue was harvested for OCT cryo-embedding. Samples were embedded in a sagittal orientation in chilled OCT and immediately flash frozen at -100 °C. All tissues were then stored at -70 °C until sectioning could be completed. The samples were acclimated for 30 minutes at -12 °C before sectioning. Sectioning was performed to target regions of interest: optic lobe, pituitary gland, pineal gland, cerebellum and olfactory lobe.

### snRNAseq preparation and sequencing

Counted nuclei were prepared for microfluidic encapsulation on the 10X Chromium instrument (10x Genomics®, Pleasanton, CA) to nanoliter-scale Gel bead-in-EMulsions (GEMs). Single-nuclei libraries were generated using the GemCode Single-Cell Instrument, Single Cell 3′ Library and Gel Bead Kit v3, and Chip Kit (10x Genomics®, Pleasanton, CA) according to the manufacturer’s protocol. Before sequencing, every library was analyzed on a Bioanalyzer high sensitivity chip to ensure the expected cDNA fragment size distribution. The appropriate number of individually barcoded GEM libraries were pooled and sequenced on a NovaSeq 6000 instrument (Illumina) with 2x150bp length using these sequencing parameters: 26 bp read 1 – 8 bp index 1 (i7) – 98 bp read 2 with 200 cycles.

### snRNAseq data processing

We downloaded the annotated reference genome AstMex3_surface (Warren et al. 2024) from NCBI (accession GCF_023375975.1) and converted the GTF file into a format accepted by CellRanger using a custom python script (https://github.com/WarrenLab/cleanup-ncbi-gtf). We prepared a reference from the downloaded fasta and reformatted GTF using CellRanger v7.2.0 mkref command with default parameters. We then aligned all snRNA-seq libraries to this reference and computed raw counts matrices for each sample using the CellRanger count command with default parameters. We filtered out ambient signal from each raw counts matrix by running cellbender remove-background (Fleming et al. 2023) v0.3.0 with default parameters, resulting in a single filtered counts matrix for each sample.

We next performed a standard clustering analysis using the scanpy (Wolf et al. 2018) platform v1.9.8. Briefly, we aggregated all per-sample counts matrices together, filtering out cells with fewer than 200 or more than 4,000 detected genes, more than 10,000 counts, or more than 2% of counts assigned to ribosomal genes. We normalized and log-scaled the counts matrix, chose highly variable genes, and regressed out counts per cell. We performed principal components analysis (PCA) on the resulting scaled counts matrices and then integrated the PCA bases by sample with harmony (Korsunsky et al. 2019). Finally, we computed the nearest-neighbors graph and ran leiden clustering on this graph with resolution 1.0, visualizing the results with UMAP.

### Spatial data collection

We generated spatial transcriptomic data using the Curio Seeker Slide-seq workflow (Rodriques et al. 2019). We cryosectioned fresh-frozen tissue and placed it onto a pre-fabricated slide containing a high-density array of barcoded beads, each with a unique spatial index. After gentle fixation and histological staining, we performed high-resolution imaging to document tissue morphology. We reverse transcribed in situ messenger RNA released from the tissue bound to oligo-dT sequences on the beads, incorporating both a unique molecular identifier and the bead’s spatial barcode. Following removal of the tissue, we recovered cDNA from the beads and prepared sequencing libraries for short-read sequencing on the Illumina NovaSeq 6000 instrument.

### Spatial data processing

We first created a reference using STAR (Dobin et al. 2013) v2.6.1d command genomeGenerate with default options except for “--sjdbOverhang 50”, using the reformatted NCBI GTF annotations as input (see single nucleus methods). We then ran the Curio Seeker pipeline v2.0.0 to generate counts matrices. Resulting reads were mapped to the transcriptome, decoded to bead coordinates, and integrated with the histology image to generate a spatially resolved gene expression matrix for downstream visualization and analysis.

We filtered out background noise using the process recommended by the manufacturer: we cropped out barcodes outside of the rectangle containing tissue and removed all remaining barcodes with fewer counts than a cutoff (log10 counts < 1.9). We then divided the slide into 40 µm x 40 µm windows, discarding all barcodes in any window with fewer than 7 remaining barcodes. We repeated this process with 100 µm x 100 µm windows, using 17 as a cutoff.

We assigned labels to each remaining spatial barcode through integration with the single nuclei dataset. In the scanpy (Wolf et al. 2018) platform v1.9.8, we filtered out genes in the spatial dataset present in fewer than 3 cells, then performed normalization, scaling, highly variable gene selection, regression, and PCA as described for the nuclei data. We integrated the PCA bases of the nuclei and spatial data using harmony (Korsunsky et al. 2019) with 15 iterations. We made a nearest neighbor index of the integrated nuclei basis and looked up the 5 nearest neighbors for each spatial barcode with this index. We then examined the cluster assignments for these five nuclei and chose the most common assignment as the annotation for this spatial barcode. If all five nuclei were assigned to different clusters, we did not assign a label to the spatial barcode.

### Cluster annotation

From our snRNAseq data, we produced a list of marker genes for each cluster by performing a t-test with overestimated variance to find genes with significantly higher expression in the target cluster compared to all other clusters (absolute value of log2 fold change > 1, p < 0.01). To identify specific cell types, we implemented a manual process that used a combinatorial approach, including known cell-type specific gene markers in other species, particularly fishes with specific reference to a zebra fish brain atlas (Pandey et al. 2023) as well as comparisons to mouse and human scRNAseq databases such as ScType (Ianevski et al. 2022). Next, we assigned cell-type annotations to each UMAP cluster based on both the genes expressed in that cluster and where the clusters were located spatially in the brain.

For the clusters that expressed canonical neuron markers, such as *elavl3 and elavl4 (HuC),* we differentiated neuron type by the expression of key neurotransmitter marker genes for glutamate, GABA, acetylcholine, and the monoamines. We then further divided the glutamatergic neurons by the specific type of vesicular glutamate transporter. We also identified glial subtypes. Specific immune cell types were identified among all subclusters based on their shared gene expression patterns using a prior head kidney evaluation of immune cells in *A. mexicanus (Peuß et al. 2020)*. For instance, the *apoeb* gene is a marker expressed in microglia.

### Differential composition by cell type

We used scCODA (Büttner et al. 2021) v0.1.9 to perform all differential composition analyses with population (i.e., cave vs. surface) and sex as covariates and FDR=0.05.

### Hierarchical clustering and variable comparison

We divided cells into groups, where each group is composed of cells from the same fish assigned to the same cell type, then performed UPGMA clustering of these groups based on the first 20 principal components of the normalized and scaled cell x gene expression matrix using scanpy’s dendrogram function with method=”average”. To calculate mutual information between topology and three explanatory variables (cell type, sex, and ecotype) for each internal (i.e., non-leaf) node of the tree, we created two vectors with length equal to the number of leaves descending from that node. The first vector represented topology, with all leaves descending from the current node’s left child encoded as 0 and all leaves descending from the current node’s right child encoded as 1. The second vector represented the explanatory variable, with each category of that variable encoded as a different integer. The mutual information between topology and that node was then calculated using the scipy function normalized_mutual_info_score with the two vectors as arguments. Finally, we calculated the weighted mean of internal node depth (i.e., distance from root of tree) multiplied by the mutual information score for each explanatory variable along with 95% confidence intervals for each weighted mean, based on bootstrapping with 10,000 iterations.

### Differential expression analysis

To find genes with differential expression between cave and surface in each cluster, we used the pseudobulk method (Murphy and Skene 2022), summing counts vectors for all cells in each sample into a single set of counts per sample. We then ran pydeseq2 (Love et al. 2014; Muzellec et al. 2023) v0.4.8 with design factors sex and population (i.e., cave vs. surface), considering any gene with absolute log2 fold change > 1 and adjusted p < 0.01 to be differentially expressed between cave and surface in that cell type. Python code for this analysis is provided in the project software repository. To compare these results to randomly-chosen groupings of individuals, we randomly divided the set of all individuals into two arbitrary groups of equal numbers of individuals and reran DEG analysis on each cell type with this random grouping as the only covariate.

### Pathway enrichment

Genes associated with canonical pathways and associated disease gene sets were tested for enrichment using the Ingenuity Pathway Analysis program (Krämer et al. 2014). A p-value of overlap is calculated with the right-tailed Fisher’s exact test. For our final analyses we used adjusted p-values after performing the Benjamini-Houchberg multiple hypothesis test corrections. The z-score estimates if the directions of gene expression change match expectations of activation or inhibition of the various diseases/functions in the IPA knowledge base. These scores are a normalized deviation from random directionality agreement.

### Cell-cell communication

The inference of cell-cell communication in cavefish compared to surface fish brains was estimated using LIANA+ (Dimitrov et al. 2024). We used the previously generated *A. mexicanus* to human gene ortholog set (Roback et al. 2025) to build a LIANA database to test for enrichment of ligand-receptor interactions within and between cell types. Normalized raw counts were used as input. Interaction heat maps and dot plots were built using the outgoing or incoming signals as curated in the ligand-receptor pairs database of CellPhoneDB (Troulé et al. 2025). Only interactions with p ≤ 0.05 were used to generate the resulting interaction heat map and dot plots. To estimate the multi-dimensional relationships of sender and receiver cells in the cave compared to surface fish we used the Tensor-cell2cell method (Armingol et al. 2022). Code used to perform these analyses is available in the project code repository.

## Additional information

## Acknowledgements

The authors thank Laurent Frantz, William Jeffery, Johanna Kowalko, and Max Shafer for discussions regarding this project; Carolina Zertuche-Mery, Aakriti Rastogi, Ansa Cobham, Jasmin Camacho, Anoja Perara, and Ana Santacruz for helping to arrange samples and assist with dissections; Seth Malloy for technical support with spatial tissue preparation; Nathan Bivens, Grant Zane, Natalia Karasseva, Skyler Kramer, and Lyndon Coghill in the University of Missouri Genomics Technology Core; Melany Madrid for bioinformatic discussions; and Evan Lloyd for contributions to cell annotations. The Stowers Institute Aquatics Team performed animal care. This work was funded by the National Institutes of Health, grant R24 OD030214 to WCW, ACK, and NR. Computation for this work was performed on the high performance computing infrastructure provided by Research Computing Support Services and in part by the National Science Foundation under grant number CNS-1429294 at the University of Missouri, Columbia MO. The authors gratefully acknowledge the Leibniz Supercomputing Centre for funding this project by providing computing time on its Linux-Cluster.

## Author contributions

This project was conceived and led by WCW, ACK, and NR. RAC, MX, and KG performed dissections and single-cell library preparation. ESR, KG, MX, RP, RAC, SK, YI, and KM analyzed single-nuclei and spatial RNA-seq data. ESR, KG, WCW, and YI wrote the first draft of the manuscript. All co-authors edited and approved the final draft of the manuscript.

## Ethics statement

All *A. mexicanus* tissue samples were processed from fishes reared at the Stowers Institute aquatic facility under IACUC approved protocol (Protocol ID: 2021-126).

## Data availability

All raw sequence data and count matrices generated in this study are available in the Gene Expression Omnibus under accession GSE335564. Processed and annotated cell data are available on the UCSC Cell Browser at https://astyanax-adult-brain.cells.ucsc.edu/.

## Code availability

Scripts and notebooks used to process the data in this study are available on GitHub (https://github.com/WarrenLab/astyanax-adult-brain-atlas) and with a permanent DOI at [TBA].

## Supplementary Figure Captions

**Supplementary Figure 1**. UMAP of full snRNAseq dataset, with clusters labeled by number according to size, with 0 the largest cluster and 35 the smallest cluster.

**Supplementary Figure 2**. Spatial barcodes assigned to each snRNA cluster.

**Supplementary Figure 3**. Marker gene expression for neurons (**a**) and glia (**b**).

**Supplementary Figure 4**. Cell type composition by sex. Clusters for which composition is different between male and female samples at FDR = 0.1 are denoted by *. 95% confidence intervals are shown.

**Supplementary Figure 5**. UMAP of immune subclusters, with clusters labeled by number according to size.

**Supplementary Figure 6.** Marker gene expression for subclustered immune cells.

**Supplementary Figure 7.** Cell type composition for immune cells by sex. Clusters for which compositions is different between male and female samples at FDR = 0.05 are denoted by *. 95% confidence intervals are shown.

## Supplementary Table Captions

**Supplementary Table 1.** Cell type labels, spatial locations, and marker genes for 35 initial brain clusters.

**Supplementary Table 2.** Genes expressed more in each cluster than all other clusters at multiple hypothesis adjusted p < 0.01. “p_adj” is the multiple hypothesis adjusted p-value; LFC is the log2-scaled fold change of (expression of gene in given cluster / expression of gene in all other clusters); “pct.1” is the percentage of cells in this cluster expressing the gene; “pct.2” is the percentage of cells in all clusters expressing this gene.

**Supplementary Table 3.** Cell type composition changes for covariates ecotype and sex at FDR = 0.05.

**Supplementary Table 4.** Differentially expressed genes between cave and surface by cell type (LFC > 0 indicates higher expression in surface).

**Supplementary Table 5.** Differentially expressed genes between male and female by cell type (LFC > 0 indicates higher expression in male).

**Supplementary Table 6.** Pathways enriched for differentially expressed genes between cave and surface.

**Supplementary Table 7**. Genes expressed more in each immune subcluster than all other immune subclusters at multiple hypothesis adjusted p < 0.01. “p_adj” is the multiple hypothesis adjusted p-value; LFC is the log2-scaled fold change of (expression of gene in given cluster / expression of gene in all other clusters); “pct.1” is the percentage of cells in this cluster expressing the gene; “pct.2” is the percentage of cells in all clusters expressing this gene.

**Supplementary Table 8**. Cell type labels and marker genes for immune subclusters.

**Supplementary Table 9**. Cell type composition changes among immune subclusters for covariates ecotype and sex at FDR = 0.05.

**Supplementary Table 10.** Differentially expressed genes between cave and surface by cell type, immune subclustering (LFC > 0 indicates higher expression in surface).

**Supplementary Table 11.** Differentially expressed genes between male and female by cell type, immune subclustering (LFC > 0 indicates higher expression in male).

**Supplementary Table 12.** Signalling pathways enriched for differentially expressed genes between cave and surface fish, immune subclustering.

