## Supplementary figures and images for "Cave adaptation drives coordinated transcriptional remodeling across diverse cell types in the brain of a teleost fish"

### Supplementary Figure 1

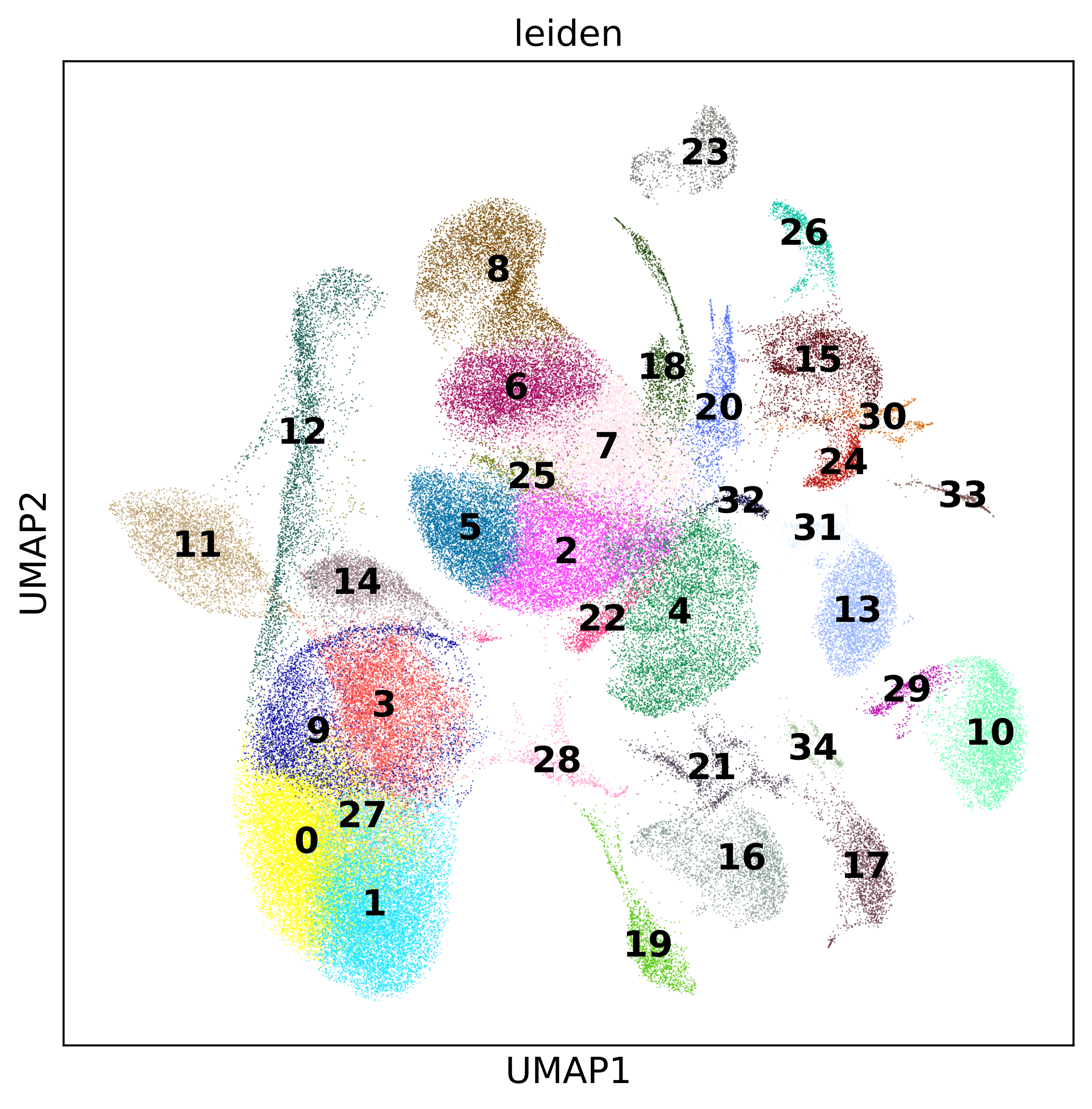

### Supplementary Figure 2

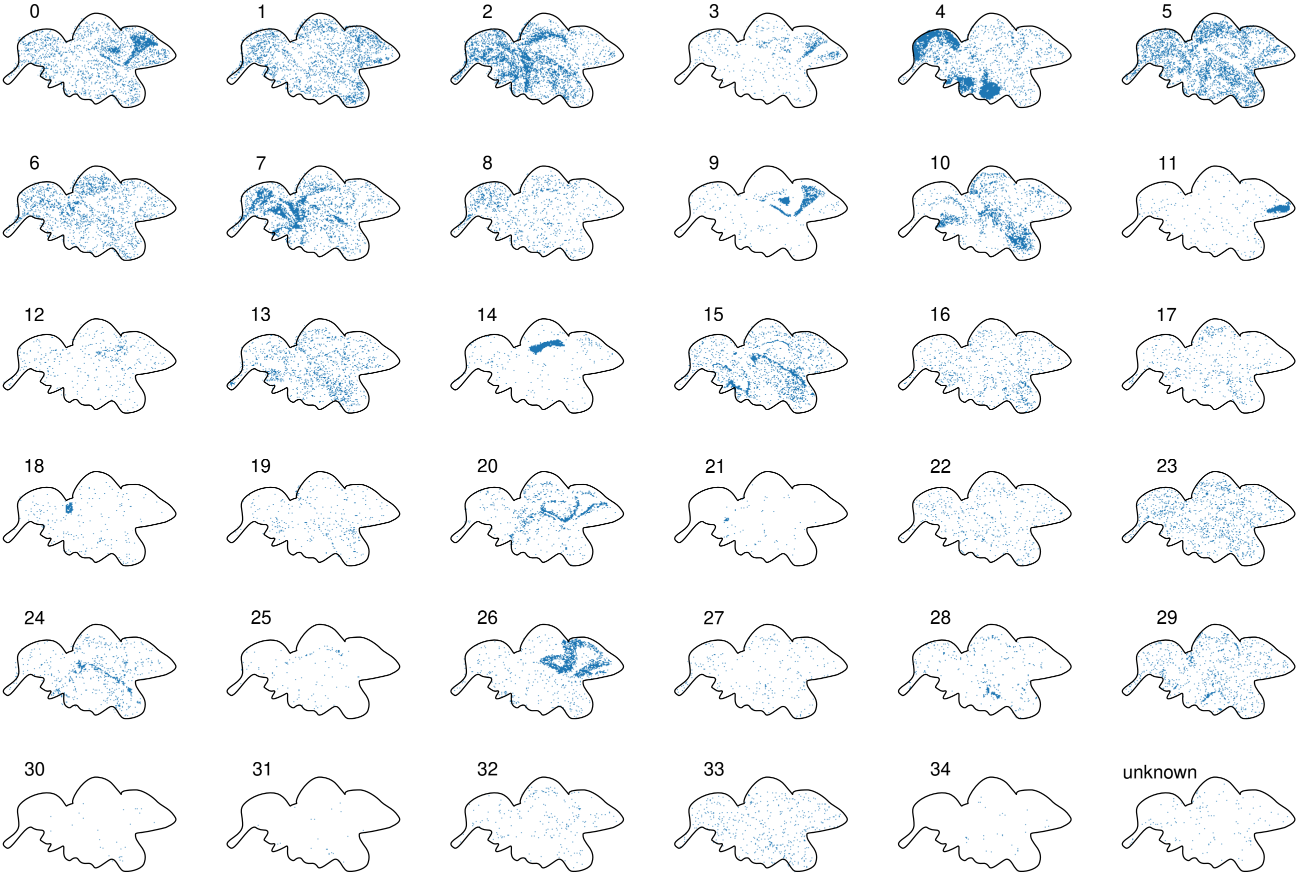

### Supplementary Figure 3

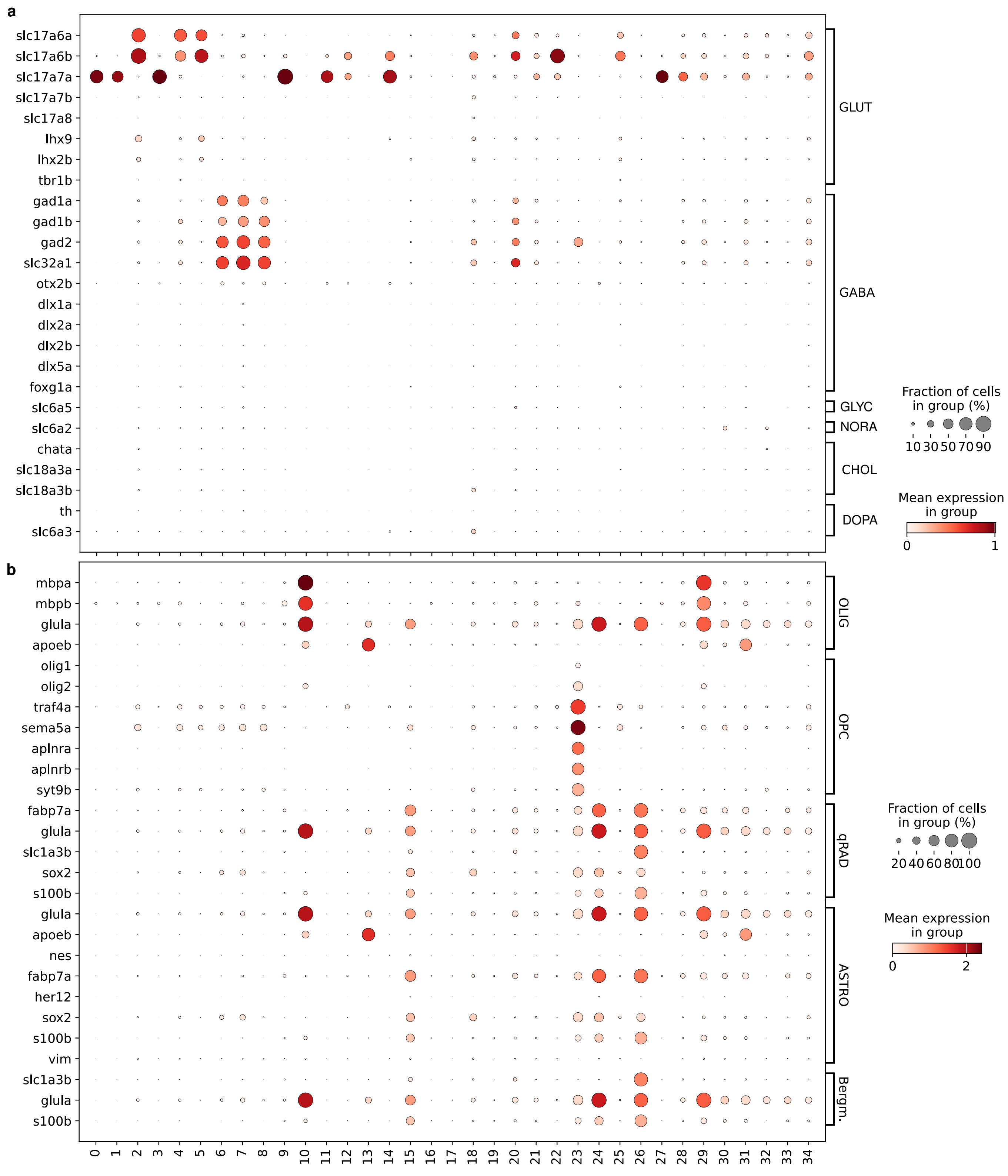

### Supplementary Figure 4

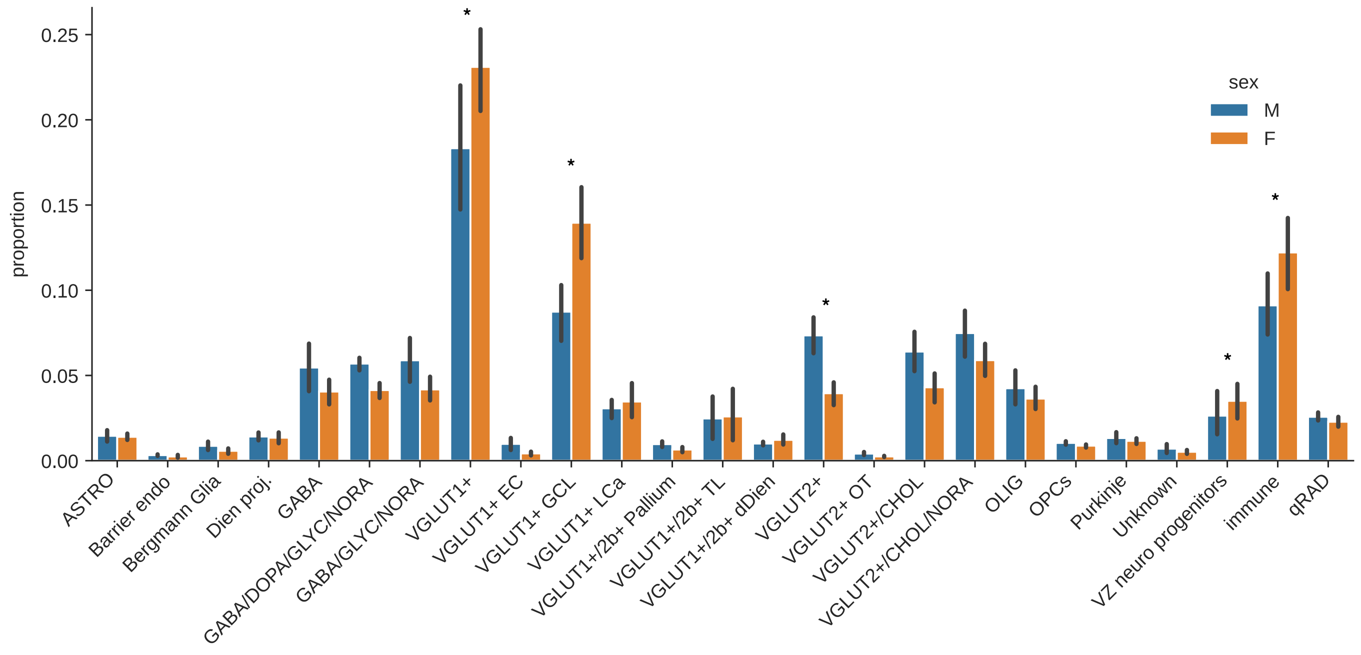

### Supplementary Figure 5

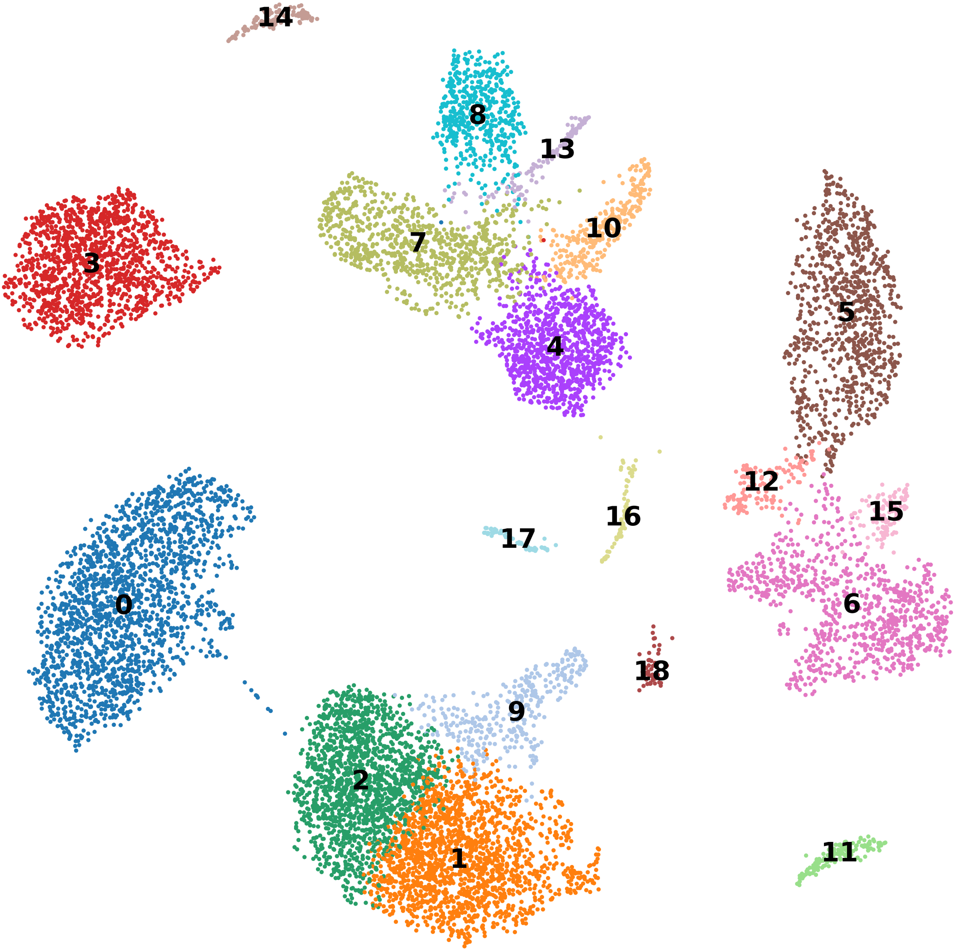

### Supplementary Figure 7

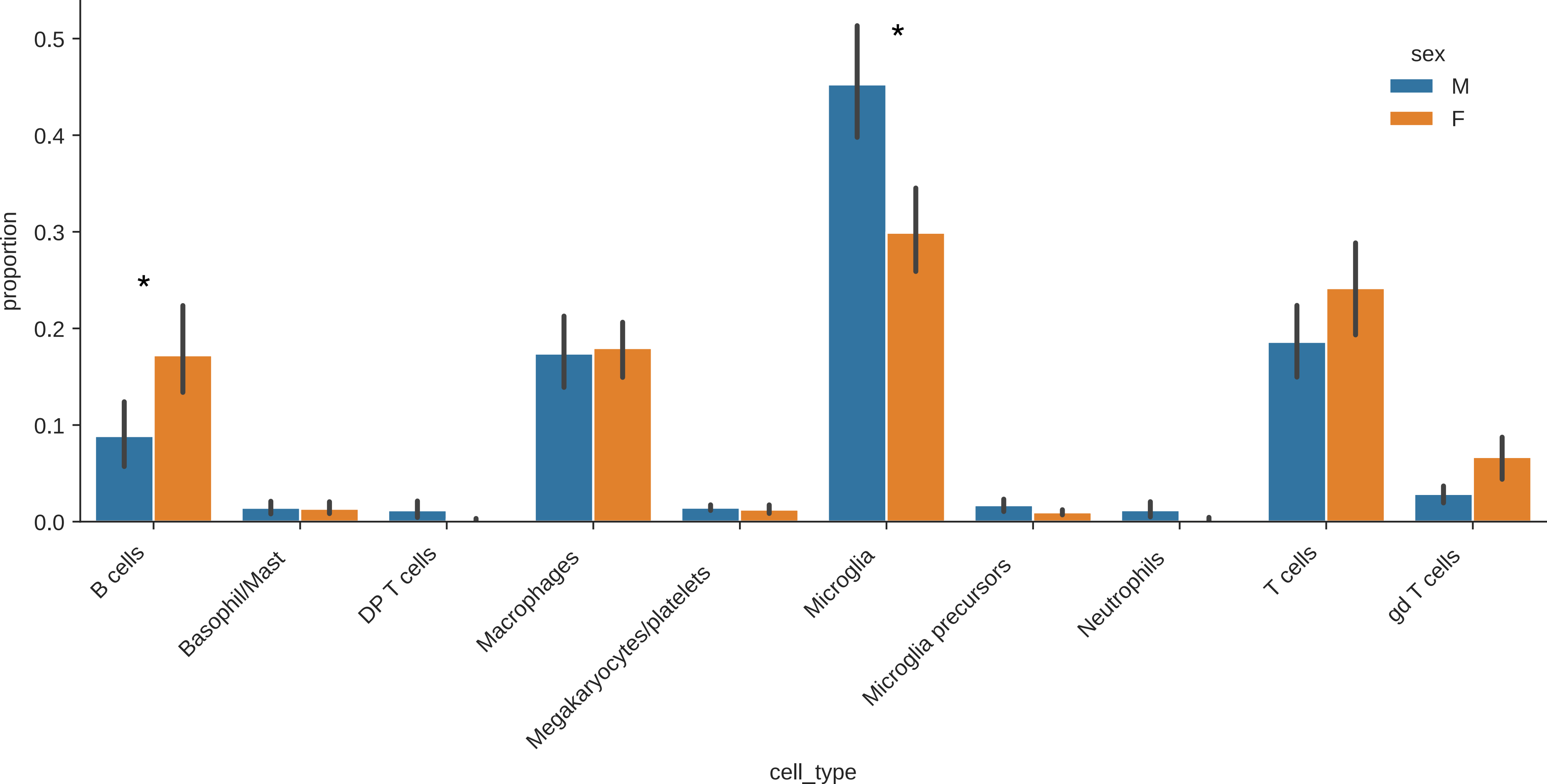

### Supplementary Table 6

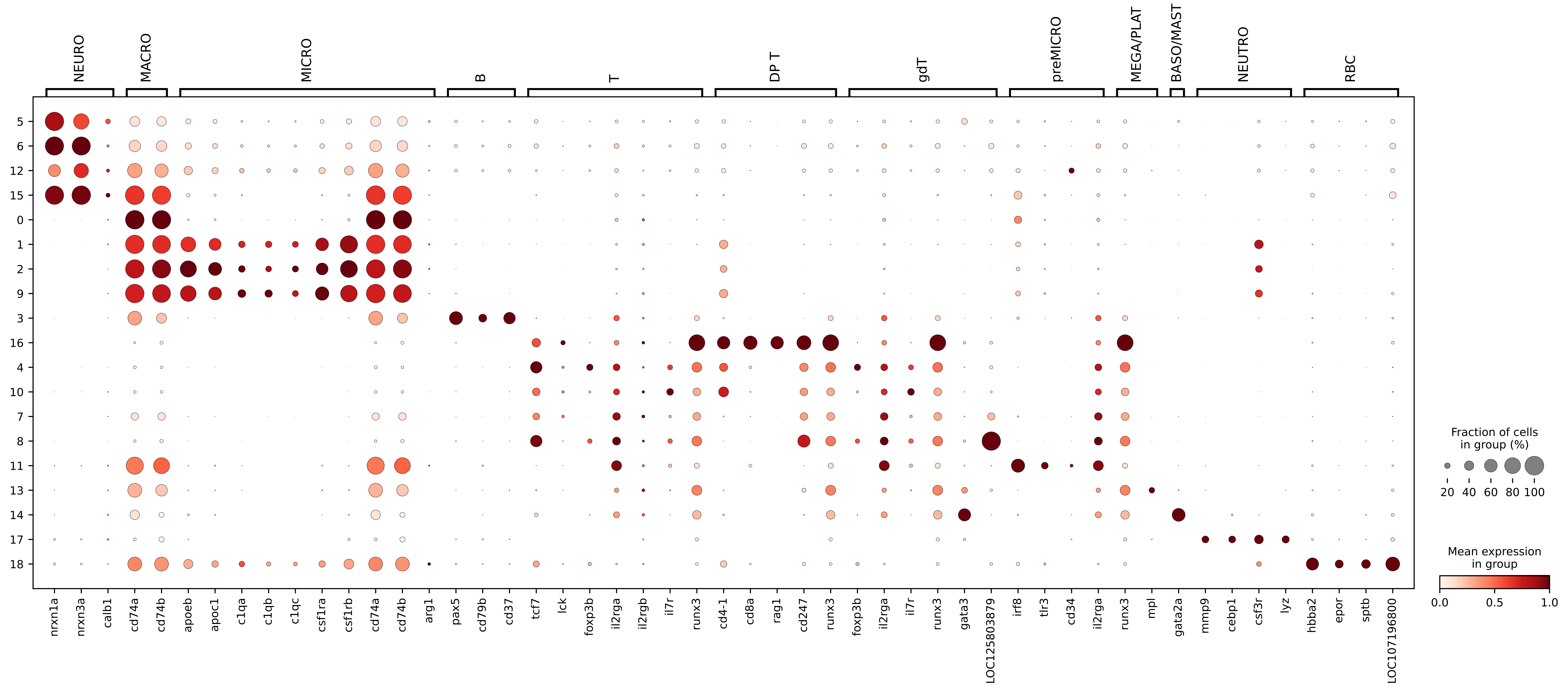
